# REverse transcriptase ACTivity (REACT) assay for point-of-care measurement of established and emerging antiretrovirals for HIV treatment and prevention

**DOI:** 10.1101/2024.08.13.607804

**Authors:** Cara Brainerd, Maya A. Singh, John Tatka, Cosette Craig, Shane Gilligan-Steinberg, Nuttada Panpradist, Megan M. Chang, Barry Lutz, Ayokunle O. Olanrewaju

**Author notes:** **Corresponding author:** Ayokunle Olanrewaju.

## Abstract

Maintaining adequate levels of antiretroviral (ARV) medications is crucial for the efficacy of HIV treatment and prevention regimens. Monitoring ARV levels can predict or prevent adverse health outcomes like treatment failure or drug resistance. However, conventional ARV measurement using liquid chromatography-tandem mass spectrometry (LC-MS/MS) is slow, expensive, and centralized delaying clinical and behavioral interventions. We previously developed a rapid enzymatic assay for measuring nucleotide reverse transcriptase inhibitors (NRTIs) – the backbone of HIV treatment and prevention regimens – based on the drugs’ termination of DNA synthesis by HIV reverse transcriptase (RT) enzyme. Here we expand our work to include non-nucleoside reverse transcriptase inhibitors (NNRTIs) – an ARV class used in established and emerging HIV treatment and prevention regimens. We demonstrate that the REverse Transcriptase ACTivity (REACT) assay can detect NNRTIs including medications used in oral and long-acting/extended-release HIV treatment and prevention. We demonstrate that REACT can measure NNRTIs spiked in either buffer or diluted plasma, and that fluorescence can be measured on both a traditional plate reader and an inexpensive portable reader that can be deployed in point-of-care (POC) settings. REACT measured clinically relevant concentrations of five NNRTI spiked in aqueous buffer. REACT measurements showed excellent agreement between the plate reader and the portable reader, with a high correlation in both aqueous buffer (Pearson’s r=0.9807, P < 0.0001) and diluted plasma (Pearson’s r = 0.9681, P < 0.0001). REACT has the potential to provide rapid measurement of NNRTIs in POC settings and may help to improve HIV treatment and prevention outcomes.

## Introduction

Antiretroviral (ARV) therapy and pre-exposure prophylaxis (PrEP) are cornerstones of HIV treatment and prevention, with diverse options including daily oral pills, monthly vaginal rings, and bi-monthly injections, that can be tailored to the needs of various populations. Despite advancements in ARV medications, including combination therapies and long-acting formulations, maintaining sufficient drug levels remains crucial for ARV efficacy.(1–3) Insufficient drug levels arise for several reasons including inconsistent medication adherence, treatment interruptions, and interindividual pharmacokinetic variations.(4–6) Key populations in HIV treatment and prevention, including elderly and pediatric individuals, pregnant people, and those with comorbidities (e.g. tuberculosis) may have additional challenges achieving adequate drug levels.(7–10) Persistent low-level ARV exposure heightens the risk of drug resistance, endangering both individual and population-level HIV treatment and prevention outcomes.(4,11,12)

There is growing evidence that incorporating drug-level feedback (DLF) into clinical trials and routine care is associated with increased medication adherence and improved HIV-related health outcomes. For example, a retrospective cohort study in Italy found that integrating therapeutic drug monitoring into routine care for people living with HIV (PLWH) reduced hospitalization time and treatment costs.(6) More recently, the MTN-034/REACH trial, which provided DLF as part of a menu of tailored adherence interventions for adolescent girls and young women (AGYW) receiving PrEP in South Africa, Uganda and Zimbabwe, observed significantly increased adherence levels compared to previous studies.(13,14)

However, the gold standard for ARV measurement, liquid chromatography-tandem mass spectrometry (LC-MS/MS), is unsuitable for routine DLF due to its high cost, expensive infrastructure requirements, and lengthy turnaround times.(13–15) There is growing interest in developing tests for rapid and accurate HIV DLF in point-of-care (POC) settings. For example, several groups have developed lateral flow assays (LFAs) for rapid ARV measurement.(16–19) Notably, urine LFAs were developed to measure tenofovir (TFV), a nucleoside reverse transcriptase inhibitor (NRTI) used in all oral PrEP regimens and ≥ 90% of HIV treatment regimens. Recent results from the Point-of-Care Urine Monitoring of Adherence (PUMA) study showed up to 72% PrEP adherence among cisgender women in Kenya when real-time DLF was provided using urine TFV LFAs, versus 45% in the standard of care group (without any DLF) at 12 months.(20) Other studies have shown that DLF with urine TFV LFAs can indicate the likelihood of virological suppression and the risk of drug resistance among people living with HIV (PLWH).(20–22)

Our group recently developed an alternate approach for measuring ARVs based on their enzyme inhibition activity. We previously described the REverSe TRanscrIptase Chain Termination (RESTRICT) assay that measures NRTIs based on the drugs’ termination of DNA synthesis by the HIV reverse transcriptase (RT) enzyme.(23–25) RESTRICT is fast (30 min), uses readily available DNA synthesis reagents, employs blood dilution as a simple sample preparation strategy, and provides results that correlate with LC-MS/MS.(24)

However, RESTRICT has so far been limited to NRTIs, representing only one class of ARVs. Non-nucleoside reverse transcriptase inhibitors (NNRTIs) are a well-established ARV class that have been used for decades in HIV treatment regimens. Although NNRTIs were recently replaced in first-line regimens by integrase strand transfer inhibitors (INSTIs), they still play a critical role in the portfolio of HIV treatment and prevention drugs. For example, the NNRTI rilpivirine (RPV) is included in recently approved bi-monthly long-acting injectable HIV treatments,(26) while dapivirine (DPV) is the sole drug in monthly extended-release vaginal rings for HIV prevention among AGYW.(27)

Although these new treatment and prevention regimens containing NNRTIs are intended to be long-acting, there is still a strong need for DLF in routine care and clinical trials. For example, AGYW receiving monthly vaginal rings in the MTN-034/REACH trial expressed a desire for real-time DLF and their healthcare providers concurred that it helped to facilitate candid conversations about the underlying reasons for non-adherence.(14) Meanwhile, therapeutic monitoring of long-acting injectable RPV is recommended by the French National Agency for Research on AIDS (ANRS MEI) to minimize the risk of treatment failure in special circumstances including missed or delayed injections, high body mass index (BMI) ≥30 kg/m^2^, known NNRTI resistance viral subtype (A1/A6), or concurrent drug regimens known to alter RPV plasma concentrations.(28)

In this study, we extend the RESTRICT assay from NRTIs to NNRTIs. NNRTIs inhibit DNA synthesis by binding to an allosteric site on the RT enzyme, inducing a conformational change that inhibits its DNA synthesis.(29) We have renamed this expanded assay, that measures both NRTIs and NNRTIs, the **RE**verse transcriptase **ACT**ivity (REACT) assay to reflect that it can operate even in the absence of DNA chain termination. We demonstrate proof-of-concept measurement of five NNRTIs in buffer solutions. We also show that REACT can measure NNRTIs spiked into human plasma. Finally, we demonstrate the feasibility of performing REACT in a portable reader, which could enable POC HIV DLF in both clinical trials and routine care.

## Materials and Methods

The following reagents were obtained through the NIH HIV Reagent Program, NIAID, NIH: etravirine [ETV] (HRP-11609, contributed by DAIDS/NIAID), rilpivirine [RPV] (HRP-12147, contributed by the NIH HIV Reagent Program), nevirapine [NVP] (HRP-4666, contributed by the NIH HIV Reagent Program). Doravirine [DOR] (B2693-475314, BOC Bioscience), dapivirine [DPV] (Adooq Bioscience, A12587), etravirine, rilpivirine, and nevirapine were reconstituted in dimethyl sulfoxide (DMSO) (Fisher Scientific, BP231-100). NNRTI concentrations were tested in serial dilutions spanning 8 orders of magnitude (10^-3^ M to 10^-10^ M), except NVP which required higher concentrations (5×10^-2^ M to 10^-8^ M) to produce a full sigmoidal curve. We used the same DNA template and primer as in our past work measuring NRTIs.(23)

REACT assays were performed in an RT buffer containing 60 mM Tris (Invitrogen, AM9855G), 30 mM KCl, (Sigma Aldrich, 101553), 8 mM MgCl_2_ (Sigma Aldrich, 63069), and 10 mM dithiothreitol (Sigma Aldrich, 20-265) adjusted to a pH 8.0 with 1 M NaOH (Sigma Aldrich, S8263). REACT assays were performed as previously described (23–25), but with final concentrations of 1000 nM dNTPs, 10 nM template, and 100 nM primer. Each assay required 20 µL of Master Mix, 10 µL of NNRTI in DMSO, and 10 µL of HIV-1 RT (Worthington Biochemical, LS05009). Reactions were performed in black, non-binding, flat-bottom polystyrene 96-well plates (Corning Incorporated, 3650). 10 µL of RT (at a final concentration of 0.053 units/mL per reaction) was added as the last reagent to initiate DNA synthesis prior to transferring the 96-well plates to a plate reader (SpectraMax iD3, Molecular Devices) for incubation at 37°C for 30 minutes with medium shaking (517 rpm). Subsequently, 40 µL of Quant-iT™ PicoGreen™ [PG] (Invitrogen, P7581) intercalating dye, diluted 1:200 in 1xTE buffer (Integrated DNA Technologies, 11-05-01-09), was added to quench the reaction and provide fluorescence readout. After an additional 1-minute incubation with medium shaking to mix PG with the master mix, readout was performed on a plate reader.

“No enzyme” negative controls containing 10 µL of RT buffer instead of the RT enzyme and “no drug” positive controls containing 10 µL of DMSO instead of a drug dilution were used to normalize fluorescence readings. Normalized fluorescence values were fitted to four-parameter logistic regression curves using GraphPad Prism Software (GraphPad Software Inc.).

### REACT assays in portable reader

REACT assays in the Harmony portable reader (30) were performed as described above with DPV in DMSO spanning 8 orders of magnitude in concentration (10^-3^ M to 10^-10^ M). Mechanical, optical, and electrical components of the Harmony reader were assembled as previously described.(30) After fluorescence readout on the plate reader, 60 µL of each sample was transferred from the 96-well plate to individual PCR tubes (Millipore Sigma, BR781320) and placed in the portable reader. Each sample was measured in each of the four available wells for 25 seconds, with a fluorescence measurement taken every 1.2 seconds. The average of these values during the 25-second measurement was taken as the representative fluorescence value to reduce the impact of background noise on the reading. Fluorescence readout was normalized to “no enzyme” and “no drug” controls and fit to a four-parameter logistic regression curve. The resultant fluorescence values from the portable reader were compared to measurements from the plate reader by performing a simple linear regression and calculating the Pearson correlation coefficient (GraphPad Prism).

### REACT with spiked plasma samples

Gender-pooled, unfiltered human plasma from healthy participants collected into K2 EDTA anticoagulant was acquired from BIOIVT (HUMANPLK2-0000283) and diluted 1:4 in nuclease-free water (Invitrogen, AM9937). Plasma was spiked with DOR dilutions in DMSO for final DOR concentrations spanning 8 orders of magnitude matching the DOR-spiked buffer concentrations (10^-3^ M to 10^-10^ M). To account for the 76% protein binding of DOR in plasma, the spiked sample concentration was multiplied by 0.24.(31) REACT was performed by substituting 10 µL of DOR-spiked plasma for the 10 µL of DOR in DMSO in the reaction. After readout on the plate reader, each sample was transferred to PCR tubes for measurement in the portable reader as described above. “No enzyme” controls and “no drug” controls were performed as described above, but with 2.5 µL DMSO and 7.5 µL of diluted plasma instead of the DOR-spiked plasma sample. Fluorescence readouts were normalized to “No enzyme” and “No drug” controls and fit to a four-parameter logistic regression curve.

## Results and Discussion

REACT measures the dsDNA synthesis activity of HIV RT enzyme as a function of the concentration of RT inhibitors present (**Fig. 1**). At high drug concentrations, DNA synthesis is inhibited leading to a low amount of dsDNA product. Conversely, low drug concentrations allow for more dsDNA synthesis. REACT uses an intercalating dye (PicoGreen™) to quantify the amount of dsDNA synthesized by providing fluorescence output that correlates with drug levels.

**Fig. 1.**
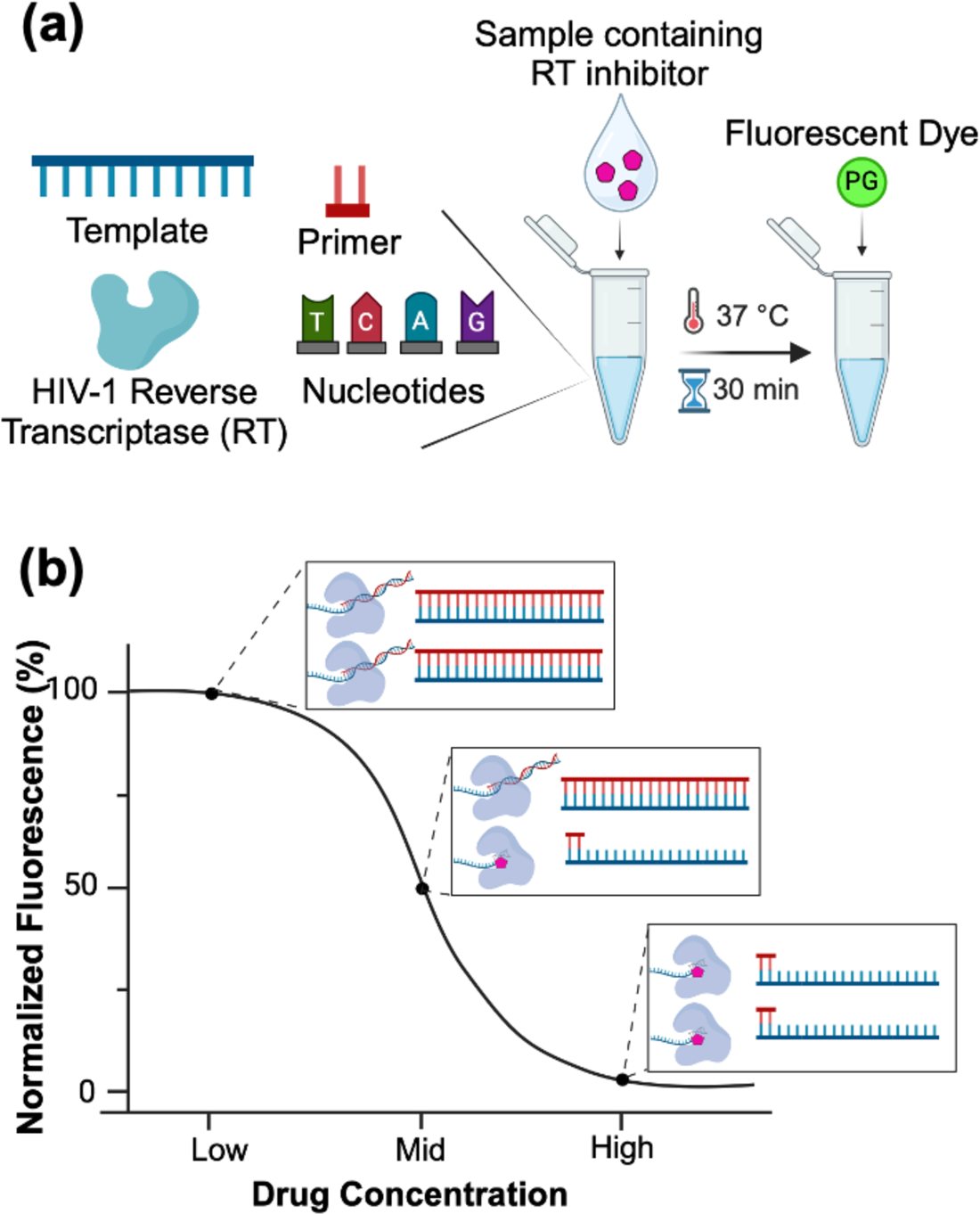
Overview of REverse transcriptase ACTivity (REACT) assay for rapid HIV drug level feedback. **(a)** REACT measures HIV reverse transcriptase (RT) inhibitors in a sample by incubating it with DNA synthesis reagents (nucleic acid templates, primers, nucleotides, and recombinant HIV RT enzyme). Following incubation at 37°C for 30 minutes, the reaction is quenched by adding PicoGreen™ intercalating dye. **(b)** Fluorescence readout measures double-stranded DNA (dsDNA) synthesis by RT as a function of drug concentration

### REACT measures multiple NNRTIs used in HIV treatment and prevention

We performed a proof-of-concept demonstration of REACT using five NNRTIs included in current HIV prevention and treatment regimens, listed in **Table 1**. We used maximum concentration (C_max_) values in pharmacokinetic trials as benchmarks for clinically relevant concentrations of each drug. REACT provided the anticipated sigmoidal curve, indicative of enzymatic inhibition reactions, across all five NNRTIs tested (**Fig. 2**). Notably, there was minimal variability between replicates, with an average coefficient of variation (CV) <5%.

**Fig. 2.**
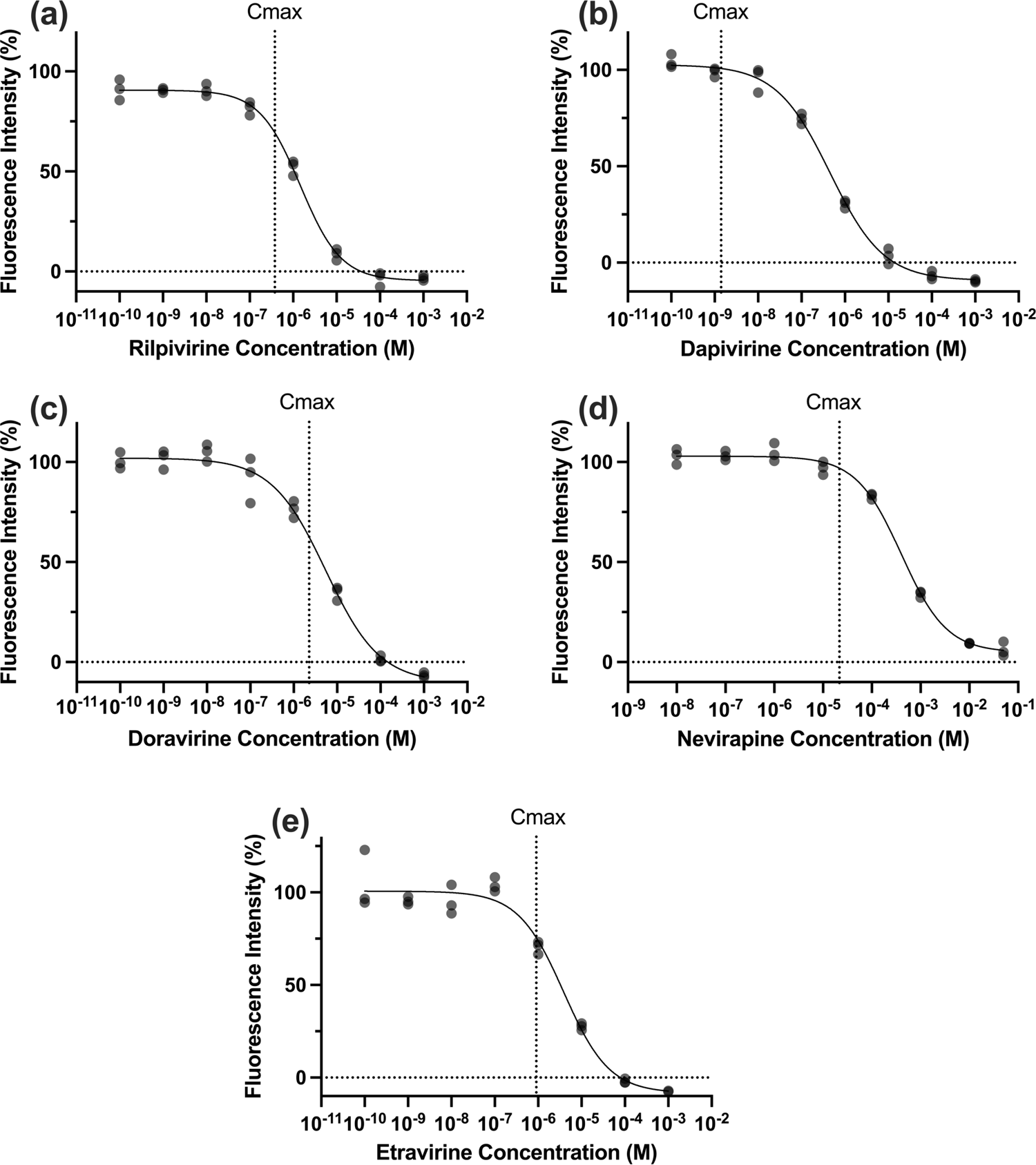
Measurement of multiple NNRTIs using REACT. Normalized fluorescence is plotted against drug concentrations of **(a)** Rilpivirine **(b)** Dapivirine **(c)** Doravirine **(d)** Nevirapine and **(e)** Etravirine. N=3. Vertical lines indicate C_max_ values in Table 1

**Table 1.**
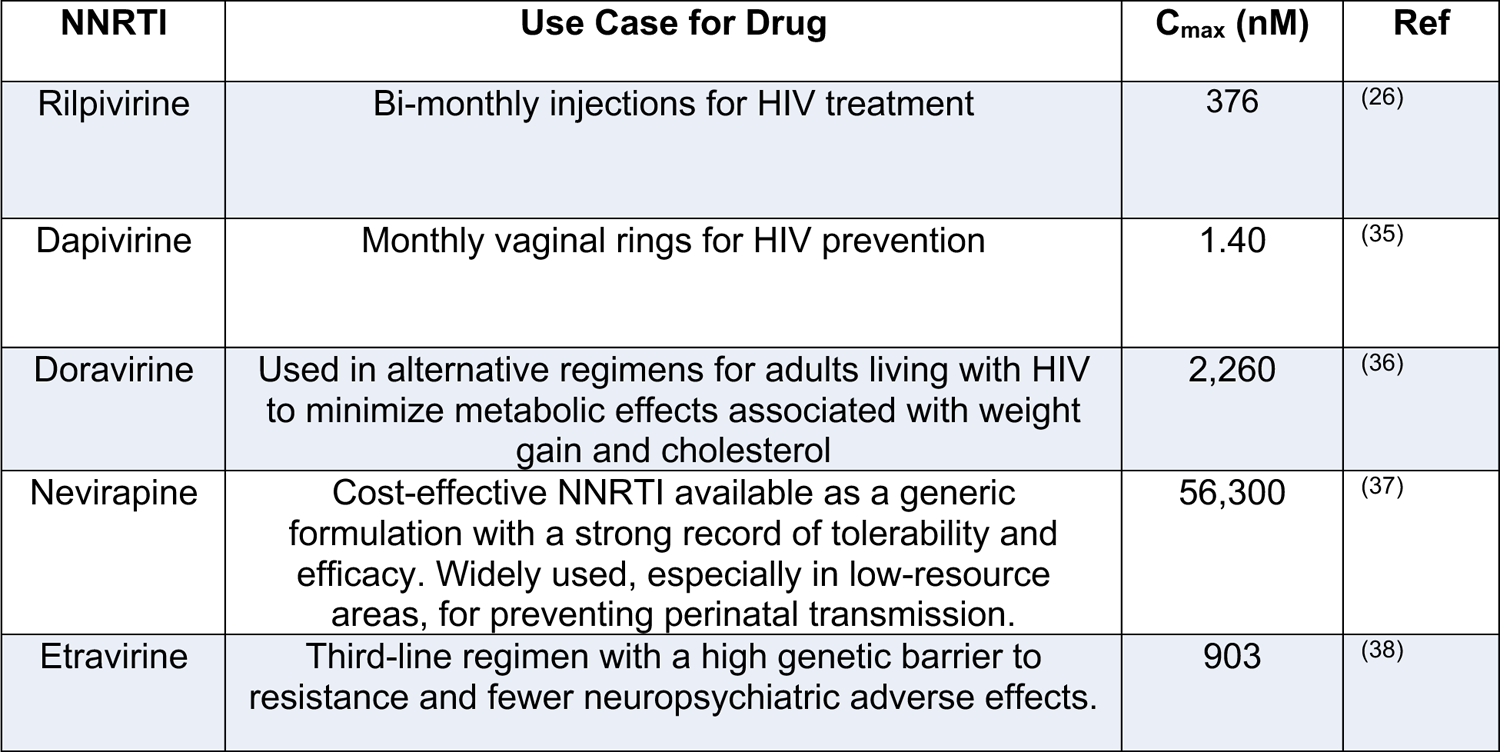
Summary of NNRTIs tested, use cases for each drug, and reported C_max_ values in plasma in pharmacokinetic studies.

| NNRTI | Use Case for Drug | $C_{\max}$ (nM) | Ref |
| --- | --- | --- | --- |
| Rilpivirine | Bi-monthly injections for HIV treatment | 376 | (26) |
| Dapivirine | Monthly vaginal rings for HIV prevention | 1.40 | (35) |
| Doravirine | Used in alternative regimens for adults living with HIV to minimize metabolic effects associated with weight gain and cholesterol | 2,260 | (36) |
| Nevirapine | Cost-effective NNRTI available as a generic formulation with a strong record of tolerability and efficacy. Widely used, especially in low-resource areas, for preventing perinatal transmission. | 56,300 | (37) |
| Etravirine | Third-line regimen with a high genetic barrier to resistance and fewer neuropsychiatric adverse effects. | 903 | (38) |

While C_max_ values can vary significantly due to inter- and intra-individual pharmacokinetic differences and testing procedures, C_max_ values are a useful metric for evaluating whether an assay provides clinically relevant results.(32–34) The linear region of REACT sigmoidal curves overlapped with the C_max_ for four of the five NNRTIs tested, illustrating the potential utility of the platform for distinguishing clinically relevant values across populations of interest.

### Integrating REACT with a Portable Reader

To demonstrate the potential of REACT as a POC test, we integrated the assay into the Harmony portable heater and fluorescence reader that was previously developed and validated for COVID-19 testing (**Fig. 3a**).(30) Harmony offers several practical advantages over traditional benchtop plate readers that make it more suitable for POC applications, particularly in terms of reduced size, weight, and cost. For example, the plate reader used for proof-of-concept REACT validation in this manuscript weighs 40 kg and occupies 3.18 m^2^ of bench space, necessitating a dedicated workstation. In contrast, Harmony weighs only 0.1 kg, representing 0.25% of the plate reader weight, and occupies just 1% of the benchtop area (0.035 m^2^) (**Fig. 3b**). Notably, Harmony costs only ∼$300 USD to assemble,(30) whereas the plate reader costs ∼$30,000 USD. While the benchtop plate reader requires a computer for data readout and analysis, Harmony results can be obtained on a cell phone. Integration into Harmony increases access to REACT especially in low-resource settings where HIV is endemic.(35)

**Fig. 3.**
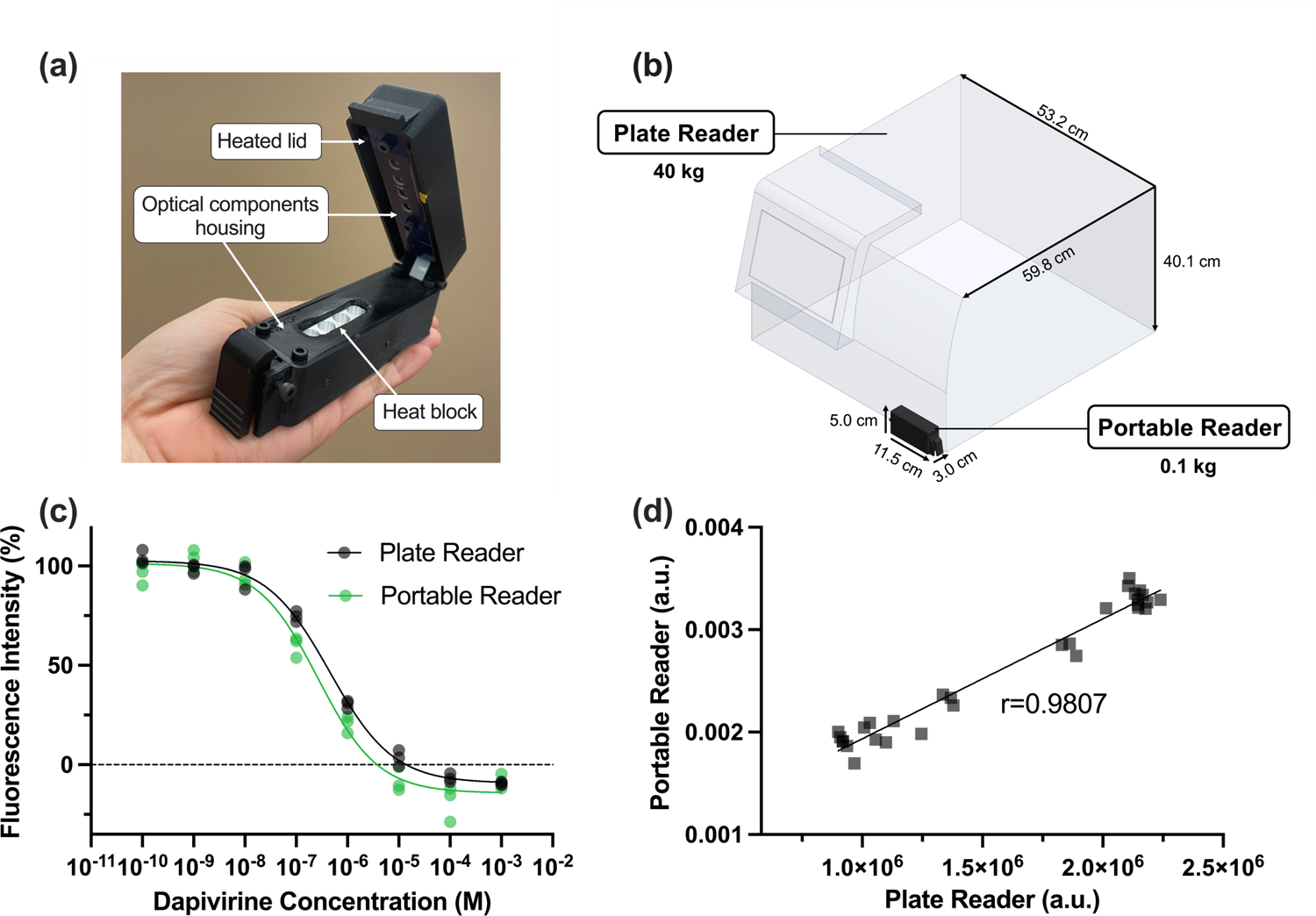
REACT results are consistent between a portable reader and a traditional benchtop plate reader. **(a)** The Harmony device consists of fluorescence detectors and a heat block with four reaction tube wells. **(b)** The Harmony portable reader offers a significant reduction in size and weight compared to a traditional plate reader. **(c)** Normalized fluorescence values indicate similar performance of REACT measurements of DPV in both readers. Plate reader data is the same as in Figure 2b. N=3. **(d)** Simple linear regression shows a correlation (Pearson’s r = 0.9807) between REACT measurements in the plate reader and measurements from a representative well in the portable reader. N=30

There was high correlation between measurements from Harmony and the plate reader (Pearson’s r=0.9807, P<0.0001) (**Fig 3c & d)**. Because Harmony only has 4 wells, we incubated REACT assays at 37°C in the plate reader during high-throughput characterization experiments before measuring fluorescence output in either Harmony or the plate reader. However, we verified that the complete incubation and fluorescence readout process on Harmony did not result in appreciable differences in fluorescence intensities (**Fig. S1**). Fluorescence output indicated in **Fig. 3c** was derived from a representative well in Harmony (see **Fig. S2** for fluorescence from all Harmony wells). Taken together, **Fig. 3** shows that REACT can be performed in a portable reader that offers substantial reductions in size, weight, and cost relative to a traditional plate reader.

### REACT with Spiked Plasma Samples

To evaluate the performance of REACT in a more complex sample matrix, we spiked doravirine (DOR) into human plasma obtained from healthy volunteers not receiving any ARVs. As in our past work, we diluted samples in water (25% v/v) as a simple sample preparation method to reduce non-specific HIV RT inhibition. **Fig. 4a** compares REACT with DOR spiked into aqueous buffer or human plasma. The x-axis of the plasma curve was adjusted to account for the 76% protein binding of DOR in plasma.(31) Raw REACT fluorescence values were higher in plasma than in buffer, likely due to autofluorescence of endogenous proteins and other plasma components.(39) Normalizing fluorescence corrected this baseline shift and showed similar dose-response curves in plasma and in buffer (**Fig. 4b**), suggesting minimal non-specific assay inhibition in diluted plasma samples. REACT assays in Harmony with DOR spiked in plasma showed excellent agreement (Pearson’s r=0.968, P<0.0001) with assays in the plate reader (**Fig. 4c & d**). Taken together, **Fig. 4** shows that REACT can measure DOR spiked into human plasma at clinically relevant concentrations using either a plate reader or portable reader.

**Fig. 4.**
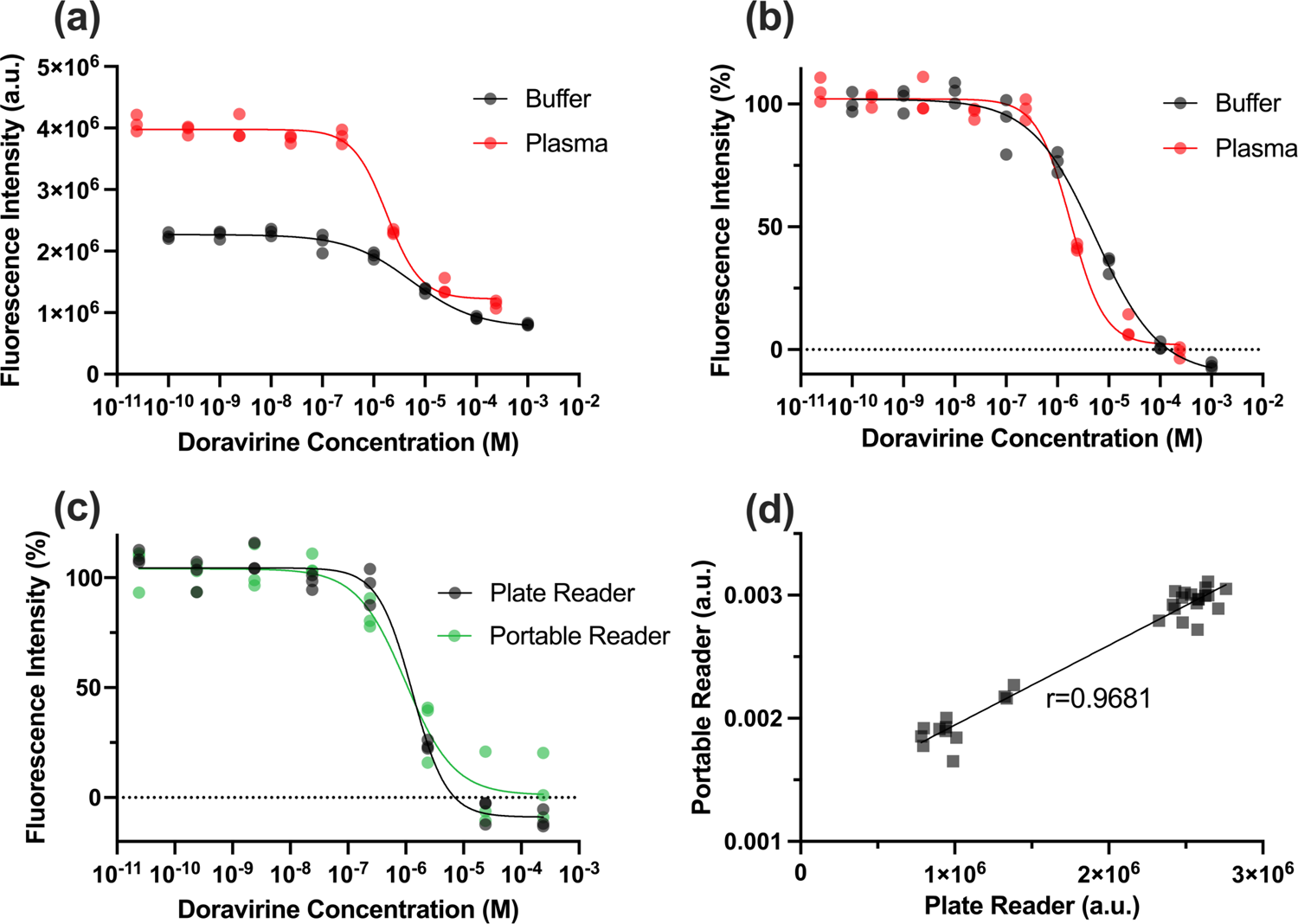
REACT provides accurate results with spiked human plasma samples. **(a)** Raw REACT fluorescence from DOR spiked into aqueous buffer and diluted (25% v/v) human plasma shows increased background fluorescence in plasma samples. The buffer curve is the same as shown in Figure 2c. **(b)** Normalized fluorescence shows excellent agreement between REACT results in spiked buffer and plasma samples. **(c)** REACT results in plasma were comparable between the plate reader and the Harmony portable reader. **(d)** Simple linear regression shows high correlation (Pearson’s r=0.9681, P<0.0001) between REACT measurements in the plate reader and measurements from a representative well in the portable reader with plasma samples. N=30. Drug concentrations in REACT assays in plasma samples were adjusted for the 76% protein binding of DOR

### Limitations of this work and future directions

Although our results show excellent performance of REACT with a complex biological matrix and a portable reader, there are still areas for improvement. One key limitation of our demonstrations so far is that we have not yet tested clinical samples containing NNRTIs and compared REACT with gold standard LC-MS/MS measurements. Testing clinical samples would allow us to evaluate the ability of the assay to differentiate between people with different ARV levels and assess correlations with health outcomes like HIV infection, viral rebound, or drug resistance.

In addition, although we have expanded from RESTRICT (which focused solely on NRTIs) to REACT (which includes NNRTIs), we have not yet included other emerging RT inhibitors like islatravir – the nucleotide transcriptase translocation inhibitor that is currently in clinical trials.(40–42) Future work will focus on emerging RT inhibitors as well as optimizing the assay to measure established drugs like tenofovir that are being integrated into long-acting and extended-release formats like implants. (43,44)

## Conclusion

The REACT assay offers a versatile solution for measuring NNRTI drug levels, addressing limitations of the gold standard LC-MS/MS such as cost and centralized laboratory requirements. We have demonstrated the detection of five NNRTIs, including RPV, DPV, DOR, NVP, and ETV, including drugs used in long-acting injectable and extended-release HIV treatment and prevention regimens. Moreover, we verified the ability to rapidly measure REACT fluorescence output in a portable reader with comparable performance to that of a traditional plate reader. This supports the feasibility of replacing the large, expensive benchtop equipment required for REACT with an inexpensive portable reader to increase accessibility to rapid DLF, particularly in resource-limited settings. Finally, we validated the ability of the REACT platform to detect an NNRTI in a complex biological matrix using both the plate reader and the portable reader. This work is innovative because it expands our prior work on enzymatic assays for HIV DLF to a new class of RT inhibitor. This activity-based assay strategy could also be useful for measuring HIV drug classes beyond RT inhibitors. Future work will validate REACT with clinical samples and compare its performance against gold standard LC-MS/MS testing.

## Declarations

### CRedIT Contributions for Authors

**Cara Brainerd:** Conceptualization; Methodology; Formal Analysis; Investigation; Writing-Original Draft Preparation; Writing-Reviewing and Editing; Visualization; Project Administration. **Maya Singh:** Validation; Formal Analysis; Investigation; Visualization. **John Tatka:** Methodology; Formal Analysis; Investigation; Visualization. **Cosette Craig:** Methodology. **Megan Chang:** Writing-Reviewing and Editing. **Shane Gilligan-Steinberg:** Resources. **Nuttada Panpradist:** Resources. **Barry Lutz:** Resources. **Ayokunle Olanrewaju:** Conceptualization, Methodology, Writing-Reviewing and Editing, Funding Acquisition, Resources, Project Administration.

## Acknowledgments

The authors are grateful for funding from the Washington Research Foundation. The authors are also thankful for NIH HIV Reagent Program, NIAID, NIH for providing etravirine, rilpivirine, and nevirapine. Figure 1 was created with Biorender.com

