## Supplemental Information for "REverse transcriptase ACTivity (REACT) assay for point-of-care measurement of established and emerging antiretrovirals for HIV treatment and prevention"

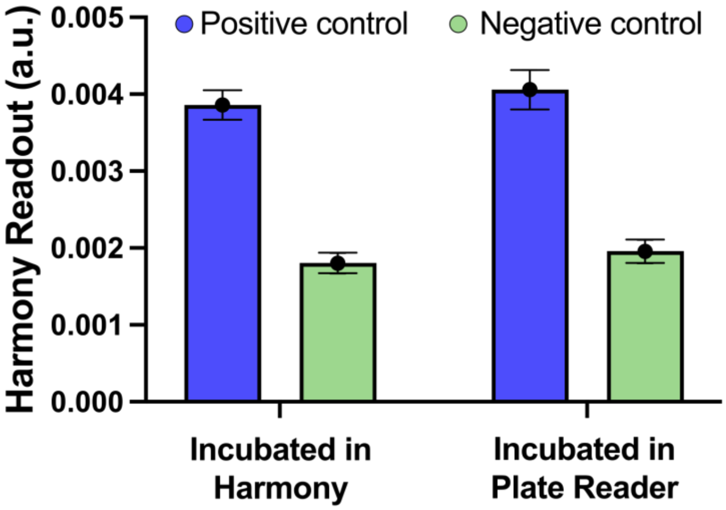


**Fig. S1 Fluorescence measurements from Harmony are not significantly affected by incubation in either the Harmony or plate reader.** Positive and Negative controls were either incubated in the Harmony or plate reader before readout of each replicate on the Harmony in all 4 wells. N=2 for Harmony incubation reactions. N=3 for plate reader incubation reactions. Error bars indicate mean and standard deviation

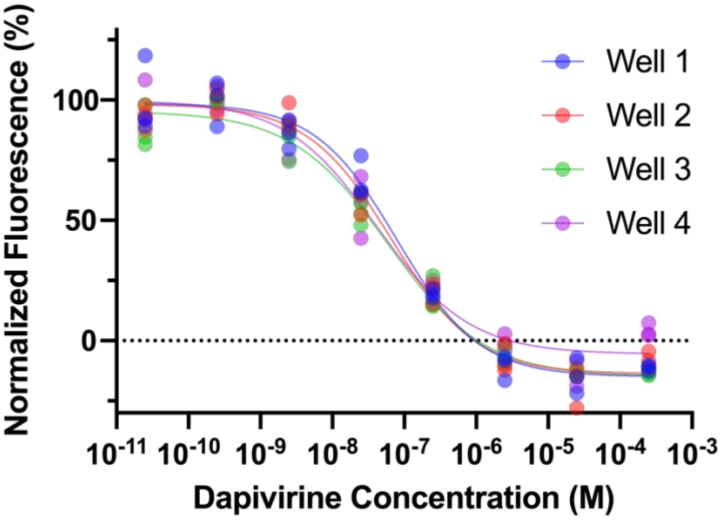


**Fig. S2 REACT detection of dapivirine in all Harmony wells**. Normalized fluorescence output of REACT is not appreciably different between wells in the Harmony. Eight drug concentrations were tested in triplicate in all 4 Harmony wells
